# Neural oscillations track the maintenance and proceduralization of novel instructions

**DOI:** 10.1101/2020.01.20.912162

**Authors:** Silvia Formica, Carlos González-García, Mehdi Senoussi, Marcel Brass

**Author notes:** Corresponding author: Silvia Formica, Ghent University, Faculty of Psychology and Educational Sciences, Department of Experimental Psychology, Henri Dunantlaan 2, B-9000 Gent.

## Abstract

Humans are capable of flexibly converting symbolic instructions into novel behaviors. Previous evidence and theoretical models suggest that the implementation of a novel instruction requires the reformatting of its declarative content into an action-oriented code optimized for the execution of the instructed behavior. While neuroimaging research focused on identifying the brain areas involved in such a process, the temporal and electrophysiological mechanisms remain poorly understood. These mechanisms, however, can provide information about the specific cognitive processes that characterize the proceduralization of information. In the present study, we recorded EEG activity while we asked participants to either simply maintain declaratively the content of novel S-R mappings or to proactively prepare for their implementation. By means of time-frequency analyses, we isolated the oscillatory features specific to the proceduralization of instructions. Implementation of the instructed mappings elicited stronger theta activity over frontal electrodes and suppression in mu and beta activity over central electrodes. On the contrary, activity in the alpha band, which has been shown to track the attentional deployment to task-relevant items, showed no differences between tasks. Together, these results support the idea that proceduralization of information is characterized by specific component processes such as orchestrating complex task settings and configuring the motor system that are not observed when instructions are held in a declarative format.

**Highlights:**

- Frontal theta power is increased during instructions implementation
- Attentional orienting in WM is analogous across maintenance and implementation
- Instructions implementation involves motor recruitment

## 1. Introduction

One peculiar aspect of human cognitive flexibility involves the ability to convert complex symbolic instructions into novel behaviors, even in the absence of prior practice (Cole, Laurent, & Stocco, 2013). Recent experimental evidence and theoretical proposals have put forward a serial heuristic model in which instructions are first encoded and maintained in a declarative format before being transformed into an action-oriented format (i.e., procedural representation) that allows executing the instructed task (Brass, Liefooghe, Braem, & De Houwer, 2017). Neuropsychological and behavioral evidence supports a dissociation between ‘knowing’ the content of an instruction and ‘doing’ the instructed cognitive or motor action, suggesting that maintaining its declarative content is insufficient to implement the instructed task, and that additional procedural codes are needed for optimal performance (Bhandari & Duncan, 2014; Duncan, Emslie, Williams, Johnson, & Freer, 1996; Liefooghe, Wenke, & De Houwer, 2012; Milner, 1963; Wenke, Gaschler, Nattkemper, & Frensch, 2009). Such immediate reformatting seems to occur only when the instruction is prepared for implementation, and not when participants merely maintain the information in a declarative manner, without the intention to implement (Liefooghe et al., 2012).

A growing number of fMRI studies has primarily focused on revealing which brain regions support the implementation of novel task sets (Demanet et al., 2016; González-García, Arco, Palenciano, Ramírez, & Ruz, 2017; Hartstra, Kühn, Verguts, & Brass, 2011; Hartstra, Waszak, & Brass, 2012; Palenciano, González-García, Arco, & Ruz, 2019; Ruge & Wolfensteller, 2010) and, more recently, on characterizing implementation-specific neural representations (Bourguignon, Braem, Hartstra, De Houwer, & Brass, 2018; González-García, Formica, Wisniewski, & Brass, 2019; Muhle-Karbe, Duncan, De Baene, Mitchell, & Brass, 2017; Palenciano, González-García, Arco, Pessoa, & Ruz, 2019; Ruge, Schäfer, Zwosta, Mohr, & Wolfensteller, 2019). These results consistently point towards a crucial role of frontoparietal regions, and in particular of the prefrontal cortex (PFC). Similarly, they provide evidence for distinct neural mechanisms supporting implementing versus memorizing novel instructions (Brass et al., 2017). However, it is still not clear to what extent declaratively representing information and proceduralization involve processes that overlap or differ qualitatively. Crucially, investigating the oscillatory dynamics of instruction implementation would allow to decompose the specific spectral components that are involved in proceduralizing S-R mappings into action-oriented codes, and to provide insights on their temporal unfolding. This would increase our understanding of the functional significance of such processes, complementing the knowledge on localization and representational format provided by fMRI research.

Specifically, we reasoned that both declarative maintenance and proactive proceduralization of novel instructions likely engage certain shared processes, such as selective attention towards the instructed content, whereas differences in brain activity should reflect the reformatting process that is specific to the implementation of the instruction. On the one hand, changes in the attentional focus are revealed by activity in the alpha frequency range (8 − 14 Hz) (Hari & Salmelin, 1997; Klimesch, Sauseng, & Hanslmayr, 2007; Senoussi, Moreland, Busch, & Dugué, 2019; van Ede, 2017). Modulations of posterior alpha band oscillations are a well-established mechanism associated with endogenous orienting of spatial attention and suppressing competition from distractors, allowing for the enhancement of task-relevant information (Bonnefond & Jensen, 2012; Jensen & Mazaheri, 2010; Sauseng et al., 2005). A substantial body of evidence reports decrease of alpha amplitude over posterior parietal regions contralateral to the attended spatial location, indicating mechanisms of orienting attention both to salient stimuli in the perceptual space and towards internally held representations (Capilla, Schoffelen, Paterson, Thut, & Gross, 2014; Gould, Rushworth, & Nobre, 2011; Mok, Myers, Wallis, & Nobre, 2016; Myers, Walther, Wallis, Stokes, & Nobre, 2015; Poch, Campo, & Barnes, 2014; Poch, Carretie, & Campo, 2017; Rihs, Michel, & Thut, 2007, 2009; Rohenkohl & Nobre, 2011; Schneider, Mertes, & Wascher, 2016; Thut, Nietzel, Brandt, & Pascual-Leone, 2006; van Dijk, Schoffelen, Oostenveld, & Jensen, 2008; Wallis, Stokes, Cousijn, Woolrich, & Nobre, 2015). Additionally, posterior alpha power has been observed to increase with the number of items relevant after attentional selection in a WM task (Jensen, 2002), also following retrospective selection (Manza, Hau, & Leung, 2014; Poch et al., 2017; Poch, Valdivia, Capilla, Hinojosa, & Campo, 2018). Therefore, we reasoned that alpha dynamics involved in the selection of internally maintained instructions should be independent of whether the task requires their implementation or mere maintenance.

On the other hand, we hypothesized that instruction implementation relies on a specific set of processes that result in the reformatting of the declarative content into action-oriented code. Therefore, we expected some oscillatory features to be specifically associated with proceduralization. First, the proactive reformatting of the instruction into a proceduralized action-bound representation should involve the allocation of cognitive control to allow the binding of stimulus and response. Ample evidence has associated control-related top-down mechanisms with larger theta (3 - 7 Hz) amplitude over mid-frontal scalp electrodes (Cavanagh & Frank, 2014; Cohen & Donner, 2013). Recent computational models attribute to theta a central role in orchestrating long-range neural communication and in binding task-relevant information (Senoussi et al., 2020; Verbeke & Verguts, 2019; Verguts, 2017). At the same time, studies showed increased theta power for WM manipulation (Itthipuripat, Wessel, & Aron, 2013; Onton, Delorme, & Makeig, 2005). Therefore, we hypothesized stronger mid-frontal theta power during preparation for instruction implementation compared to maintenance, reflecting the proactive reformatting of the S-R mapping into a proceduralized action-bound representation. Nevertheless, behavioral evidence suggests that only a small set of mappings can be proceduralized and maintained in such a format prior to the moment of their actual implementation, hinting at a limit in the number of novel instructions that can be proceduralized at once (Liefooghe et al., 2012). As we expect theta power to track the occurrence of such reformatting, we reasoned that theta dynamics should not differ from declarative maintenance when the number of mappings to implement exceeds capacity limits.

Finally, we reasoned that proactively recoding the instruction for execution, and not merely maintaining it, should induce the preparation of the instructed motor plan (Everaert, Theeuwes, Liefooghe, & De Houwer, 2014; Meiran, Pereg, Kessler, Cole, & Braver, 2015a). Therefore, we looked at suppression in beta (15 30 Hz) and mu (8 12 Hz) frequency bands over central electrodes, oscillatory features traditionally associated with motor preparation and motor imagery (Cheyne, 2013; Pineda, 2005; Rhodes, Gaetz, Marsden, & Hall, 2018; Schneider, Barth, & Wascher, 2017; Tzagarakis, West, & Pellizzer, 2015). We expected this modulation to be larger while preparing to implement the instructions, thus when an action-bound procedural representation is available, compared to the declarative maintenance of the mappings. Again, we predicted such difference to be reduced with a higher number of instructions, due to a limit in the number of action plans that can be prepared and maintained ready simultaneously.

To test our hypotheses, we recorded EEG activity while participants performed a task that encouraged a declarative maintenance of novel S-R mappings for recognition (Memorization task), and an Implementation task, intended to prompt their proactive reformatting for execution. In both cases, after the presentation of four mappings at the beginning of the trial, a retro-cue selected a subset of them as potential targets for the subsequent recognition or execution task. We reasoned that brain activity during the Cue-Target interval (CTI) would reflect the two-steps action described in a recent model of information prioritization in WM (Myers, Stokes, & Nobre, 2017). According to Myers and colleagues (2017), attention needs to be internally oriented towards the selected items, and the prioritized representations can then be reformatted into a behavior-optimized state. In the context of instruction implementation, the task-optimized representation is a procedural, action-bound code of the S-R mapping. On the other hand, successful maintenance is optimally achieved by means of a declarative representation of stimulus and response that does not entail any action plan. Therefore, during the CTI of the two tasks, we expected analogous features associated with attentional orienting, but instances of reformatting to be larger in preparation for instruction implementation.

## 2. Materials and Methods

### 2.1 Participants

Thirty-nine participants took part in the experiment (*M*_*age*_ = 21.74, *SD* = 4.50, 33 females) and received 30 euros as compensation. Sample size was not computed a priori: we aimed for thirty-five participants, which is on the upper bound of the sample size of studies investigating similar constructs (de Vries, Van Driel, Karacaoglu, & Olivers, 2018; Schneider et al., 2017; van Ede, Chekroud, Stokes, & Nobre, 2019a). All participants had normal or corrected-to-normal vision and thirty-three reported to be right-handed. Data from two participants were discarded due to low task performance (individual mean accuracy exceeded by 2.5 standard deviations the group mean accuracy in one of the two tasks and/or accuracy in response to catch trials in one of the tasks was below 60%); data from two additional participants were discarded following visual inspection because of excessive noise in the EEG recordings, resulting in a final sample size of thirty-five participants. All participants gave their informed consent prior to the beginning of the experiment, in accordance with the Declaration of Helsinki and the protocols of Ghent University.

### 2.2 Materials

The same set of stimuli was used as in previous studies on instruction implementation (Formica, González-García, & Brass, 2020; González-García, Formica, Liefooghe, & Brass, 2020). It consisted of 1550 pictures, grouped in two macro categories: animate (non-human animals) and inanimate (vehicles and musical instruments) (Brady, Konkle, Alvarez, & Oliva, 2013; Brodeur, Guérard, & Bouras, 2014; Griffin, Holub, & Perona, 2007; Konkle, Brady, Alvarez, & Oliva, 2010). All images had their background removed, were centered in a 200×200 pixels square and were converted to grayscale. Stimuli presentation and response collection were performed using Psychopy toolbox (Pierce, 2007).

### 2.3 Experimental design

Participants performed the two tasks (from now on “Implementation task” and “Memorization task”) in a blocked fashion during one single session, and the order of the two tasks was counterbalanced between participants. The structure of the trials for the two tasks was identical up to the presentation of the target (Figure 1). Each trial started with a red fixation cross presented for 2000ms (± 100ms, jittered) signaling the inter-trial interval and allowing participants to blink if needed, followed by a white fixation cross for 250ms. Next, the encoding screen containing four S-R mappings (arranged in two rows) appeared for 5 seconds. Mappings consisted of a new image associated with a bimanual response: “index” referred to both index fingers (keys r and i) and “middle” referred to both middle fingers (keys e and o). These bilateral responses were used instead of the more traditional left and right options to avoid automatic motor activations elicited by the mere presentation of lateralized response words (Bundt, Bardi, Abrahamse, Brass, & Notebaert, 2015). Out of the four presented mappings, two contained images of animals and two images of inanimate objects. Each image-response combination was presented only once during the whole duration of the experiment. The rationale of this choice was to maximize WM demands and minimize the retrieval of practiced S-R mappings from long-term memory (Meiran, Pereg, Kessler, Cole, & Braver, 2015b). After a 750ms delay, which is considered an interval sufficiently long to discard a contribution of iconic memory (Souza & Oberauer, 2016), the retro-cue appeared and remained on the screen for 250ms. This consisted of a square presented centrally, with one, two or four (i.e., neutral retro-cue) colored corner(s)^1^. The number of selected mappings is the second crucial factor in our design, namely Load. Participants were instructed that the retro-cue signaled which mapping(s) could be probed, with 100% validity (except in the case of catch trials, see below). Importantly, when the retro-cue selected two mappings, these were always on the same side of the screen. The subsequent cue-target interval (CTI) had a jittered duration, lasting on average 1750ms (± 100ms). This interval largely exceeds the 300 − 500 ms considered necessary for retro-cues to benefit performance (Souza & Oberauer, 2016) and sufficiently long to allow time-frequency decomposition of low frequencies (Cohen, 2014b). Finally, participants were presented with the target screen, which differed depending on the task. In the Implementation task, the image of one of the cued mappings was presented centrally and participants were required to press the associated pair of keys (both index or both middle fingers). In the Memorization task, one mapping (i.e., image and associated response) was presented centrally. The image in the Memorization probe belonged to one of the selected mappings (except for catch trials, see below). Participants had to report whether the presented mapping matched one of those selected by the retro-cue, by pressing with both fingers of one hand for “yes” or both fingers of the other hand for “no”. The sides for “yes” and “no” were randomly assigned on a trial basis to ensure that participants could not prepare any response during the CTI, therefore encouraging the mere declarative maintenance of the instruction memoranda. Labels with “yes” and “no” appeared at the bottom of the target screen together with the mapping. In 50% of the trials, the target screen showed the same mapping as in the encoding (“yes” response). In the other 50% of trials, one of the selected images was presented associated with a different response with respect to encoding (“no” response). Crucially, despite the similarities in trial structure, the Memorization task relied on the declarative maintenance of the mappings as no response could be prepared, whereas the Implementation task encouraged a proactive preparation of the selected mappings for the execution of the required action plan. Both tasks included catch trials (∼25% for each task), in which the target screen displayed a different image than the four displayed in the encoding screen. In this case, participants were required to press the spacebar. This was done to discourage participants to adopt specific strategies, such as memorizing only the mappings associated with one response option. Since we were interested in the brain activity before the onset of the target, EEG recordings from catch trials were analyzed together with regular trials. For each task, participants completed 5 experimental blocks, for a total of 180 trials (60 per load, Figure 1).

**Figure 1:**
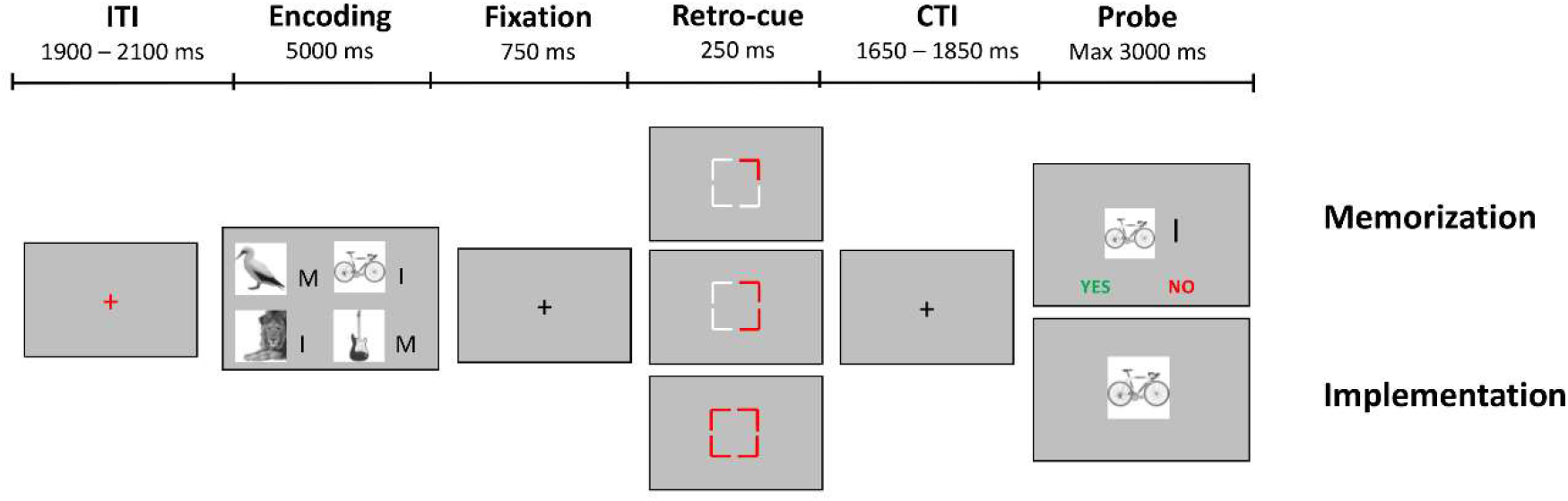
Behavioral paradigm. Participants had to first encode four S-R mappings, for subsequent recognition (Memorization task) or for a choice-reaction task (Implementation task). During the retention interval, a Retro-Cue selected one, two or all four mappings as relevant for the upcoming probe. Participants performed the two tasks separately in a block design.

Each of the two tasks was preceded by a practice session. This consisted of mini-blocks of 12 trials, including all possible load conditions (i.e. number of mappings selected by the retro-cue) and at least one catch trial. The only difference with the main task was the presence of feedback at the end of each trial, signaling the accuracy of the response or encouraging participants to respond faster in case no response was registered within the maximum response time of 3 seconds. Performance was assessed at the end of each mini-block: if accuracy was above 80%, practice was concluded, otherwise a new mini-block started, up to a maximum of 3 blocks. S-R mappings used during the practice were never presented again during the main task. The total duration of the experiment, including cap preparation, practice, main tasks and breaks was approximately 150 minutes.

### 2.4 EEG Recordings and pre-processing

Electrophysiological data were recorded using a BioSemi ActiveTwo system (BioSemi, Amsterdam, Netherlands) with 64 Ag AgCl electrodes arranged in the standard international 10 20 electrode mapping (Klem, Lüders, Jasper, & Elger, 1999), with a posterior CMS-DRL electrode pair. Two reference electrodes were positioned at the left and right mastoids. Eye movements were registered with a pair of electrodes above and below the left eye and two additional electrodes at the outer canthi of both eyes. EEG signals were recorded at a 1024 Hz sampling rate.

EEG data were preprocessed using the Fieldtrip toolbox (Oostenveld, Fries, Maris, & Schoffelen, 2010), running in MATLAB (MATLAB R2017b, The MathWorks, Inc., Natick, Massachusetts, United States). First, the data were downsampled to 512 Hz and re-referenced to the average of the mastoids. Then, a 0.5 − 45 Hz band-pass FIR filter was applied to the data, together with a Notch filter at 50 Hz and its harmonics. Data were epoched relative to the onset of the retro-cue (from −1000 to 2500 ms) and demeaned to the average of the whole epoch, to improve independent component analysis (ICA) (Groppe, Makeig, & Kutas, 2009). Only trials in which the participant performed the correct response were retained for subsequent analyses. Trials exhibiting movement artifacts or excessive noise were removed following visual inspection of the data. Next, eye movements artifacts were removed by means of ICA, using the EEGLAB (Delorme & Makeig, 2004) runica algorithm as implemented in Fieldtrip. Components to discard were selected based on their topography, the correlation between their time course and the horizontal and vertical electrooculography, and their power spectrum. For most participants, two components were removed (capturing blinks and horizontal eye movements, respectively); in six participants blinks were reflected in two components, leading to the removal of three IC, and in two participants only one component was removed. Finally, data were visually inspected again to ensure successful cleaning and one (N = 6), two (N = 2) or three (N = 1) excluded channels were interpolated by means of spherical spline interpolation (Perrin, Pernier, Bertrand, & Echallier, 1989). This cleaning procedure resulted in an average of 151.09 trials for the Implementation task (*SD* = 12.29, 83.94%) and 143.06 trials for the Memorization task (*SD* = 14.07, 79.48%). For each load condition, an average of 81.7% of trials were retained (Implementation, Load 1: 52.14 (*SD* = 3.64) trials, Load 2: 50.26 (*SD* = 5.19) trials, Load 4: 48.68 trials (*SD* = 5.74); Memorization, Load 1: 49.88 trials (*SD* = 4.84), Load 2: 47.86 trials (*SD* = 5.86), Load 4: 45.31 trials (*SD* = 5.94).

### 2.5 Spectral Analysis

For each Task x Load condition (Memorization-Load 1, Memorization-Load 2, Memorization-Load 4 and Implementation-Load 1, Implementation-Load 2, Implementation-Load 4), time-frequency analysis was performed separately using complex Morlet wavelet convolution, to estimate spectral power from 1 to 45 Hz in steps of 1 Hz. The number of cycles in the wavelet was frequency-specific, ranging from 2 at 1 Hz and linearly spaced up to 7 cycles at 45 Hz, to achieve a good trade-off between temporal and frequency precision (Cohen, 2014a). This analysis resulted in one time-frequency spectrogram for each channel, condition and participant. Condition-specific decibel normalization was then applied, using the time window between −500 and −200ms before the onset of the retro-cue as baseline. To avoid data contamination due to the smearing in time of the processing of the target, all statistical analyses are performed on the time window from 0 to 1800ms with respect to retro-cue onset, therefore leaving 100ms gap between the end of our analyses window and the earliest jittered target onset.

### 2.6 Statistical Analyses

Reaction times (RTs), error rates (ER) and averaged alpha power (see below) were separately entered in 2 (Task: Memorization vs Implementation) x 3 (Load: 1, 2, 4) repeated measure ANOVAs, performed in JASP (Jasp Team, 2019).

To evaluate the statistical significance of differences between EEG time-courses or time-frequency spectra, we adopted a cluster-based permutation approach (Maris & Oostenveld, 2007), which is appropriate to assess the reliability of neural patterns over neighboring data points. Moreover, this approach is robust against the multiple-comparison problem, as the significance of clusters found in the observed group-level data is estimated against a distribution of clusters obtained by randomly permuting the assignement of participants to each group. First, depending on the number of conditions to be compared, we performed either a two-sided t-test (two conditions) or an F-test (three conditions), at an α level of 0.05, between all data points of the observed conditions. Then, we considered as cluster a group of adjacent data points with same sign significance and as cluster-size the sum of all t- or F-values in the cluster. Next, we used 5,000 permutations of participant-level data to estimate a distribution of cluster sizes under the null hypothesis that there were no differences between conditions. The *P*-value for each cluster in the observed group-level data corresponds to the proportion of permutations in which the largest cluster size was larger than the size of the considered observed cluster. Again, we used a significance alpha level of 0.05, therefore only observed clusters whose size was larger than the size of the largest cluster in at least 95% of permutations are reported. This approach provides a statistically sound procedure to compare two conditions and establish the existence of a difference between them, reliably accounting for multiple comparisons. However, because clusters found through this procedure reflect the result of the second level inference between the observed and the null permuted distributions, no strong conclusions can be drawn with respect to their boundaries, such as the exact latency or frequency range of the effect (Sassenhagen & Draschkow, 2019).

### 2.7 Contralateral alpha suppression

Consistent findings reported alpha desynchronization over posterior regions contralateral to the attended spatial location, indicating top-down anticipatory mechanisms of attentional orienting in the perceptual or internal space (Bonnefond & Jensen, 2012; Capilla et al., 2014; Gould et al., 2011; Jensen & Mazaheri, 2010; Mok et al., 2016; Myers et al., 2015; Rihs et al., 2007, 2009; Rohenkohl & Nobre, 2011; Sauseng et al., 2005; Thut et al., 2006; van Dijk et al., 2008; Wallis et al., 2015).

To estimate the suppression of alpha oscillations contralateral to the cued hemifield, we averaged the time-frequency spectra corresponding to Load 1 and Load 2, separately for the two tasks. Trials with neutral retro-cues (Load 4) were excluded from this analysis because they did not induce attentional orienting towards one of the two visual hemifields. Based on previous literature, two pairs of electrodes were selected in right (P8, PO8) and left (P7, PO7) posterior parietal regions (de Vries, van Driel, & Olivers, 2019a; Gould et al., 2011; Schneider et al., 2017; van Ede, Chekroud, Stokes, & Nobre, 2019b). Power spectra were extracted from these electrode pairs, averaged in the alpha frequency range (8 − 14 Hz), and collapsed between retro-cues pointing to the left and right hemifield, in order to extract one power time series for the contralateral hemisphere and one for the ipsilateral hemisphere. The two time series were compared by means of cluster-based permutation testing, separately for the two tasks (see Statistics section). Additionally, the cluster-based permutation approach was also used to compare the contralateral alpha suppression between the two tasks. Namely, for each task the ipsilateral power time series was subtracted to the contralateral one, separately for each participant, and the resulting difference waves were compared. This analysis was performed to test whether the deployment of alpha oscillations to orient attention towards relevant mental representations differed between tasks.

### 2.8 Load-dependent alpha increase

Posterior alpha power has been observed to increase with the number of items relevant after attentional selection in a WM task (Fukuda, Mance, & Vogel, 2015; Jensen, 2002), also following retrospective selection (Manza et al., 2014; Poch et al., 2017, 2018). Observing such modulation in our data would be evidence in favor of an effective selection of the relevant mappings. To investigate the increase in alpha power relative to the number of retained items, we extracted the average power for the frequencies ranging from 8 to 14 Hz, separately for each condition and task, from the electrode pair Pz and POz (Jensen, 2002), across the whole time window from 0 to 1800ms with respect to the onset of the retro-cue. This operation led to one averaged alpha power value for each Task-Load combination per participant, that were entered in a 2 (Task) x 3 (Load) repeated measure ANOVA.

### 2.9 Mid-frontal theta in Implementation

The two tasks differ to the extent they require the reformatting of the encoded information in an action-oriented code. We hypothesized such transformation to rely on a top-down mechanism supported by low frequency oscillations. To test our predictions, time-frequency spectra were extracted from the CTI (0 − 1800 ms) in the frequency range 3 − 7 Hz, averaged from channels Fz and AFz. The choice of these two channels was guided by both their widespread use in the literature concerning midfrontal theta (Berger et al., 2019; Cooper et al., 2019; Onton et al., 2005; Popov et al., 2018; Senoussi et al., 2020) and a condition-independent channel selection procedure. Namely, we fitted the power spectra from all channels, computed across all trials, regardless of the Load condition and the Task, with the FOOOF toolbox (Haller et al., 2018). This method allows to decompose different components of the power spectrum (i.e., the aperiodic (1/f pattern) and periodic components) and thus to better capture the oscillatory profile of electrophysiological data. This analysis confirmed that, across conditions, channels Fz and AFz exhibited the largest peaks in the power spectra between 3 and 7 Hz (see Supplementary materials for details).

We tested the main effect of Task by comparing the time-frequency spectra of Implementation and Memorization (averaged across levels of Loads) by means of cluster-based permutation test. Similarly, we tested the main effect of Load by contrasting the spectra for each level of Load, averaged across Task condition. We then tested for a significant effect of the interaction between our two factors by running a cluster-based permutation test on the three time-frequency spectra resulting from pairwise subtraction of the two tasks for each Load level (i.e., Implementation 1 minus Memorization 1, Implementation 2 minus Memorization 2 and Implementation 4 minus Memorization 4).

### 2.10 Motor-related mu and beta suppression in Implementation

Analogously, we hypothesized that action-oriented procedural representations should entail a motor plan, reflected in stronger signatures of proactive motor preparation. We expected differences in two frequency bands associated with motor preparation, namely mu (8-12 Hz) and beta (15 − 30 Hz) (Cheyne, 2013; Pineda, 2005). To test for this, we extracted time-frequency spectra averaged across electrodes C3 and C4 from the CTI (0 −1800 ms). Next, we again used a cluster-based permutation approach outlined for mid-frontal theta, to look for significant clusters of different activity between tasks, loads, and their interaction. We selected electrodes C3 and C4 because we expected the effect to be maximal over motor and pre-motor cortices^2^ (Marchesotti, Bassolino, Serino, Bleuler, & Blanke, 2016; McFarland, Miner, Vaughan, & Wolpaw, 2000). Moreover, we averaged across the two of them because the goal of this comparison was to investigate pre-movement preparatory oscillatory activity, rather than the effector-specific M1 activation (Neuper & Pfurtscheller, 2001; Pfurtscheller, Brunner, Schlögl, & Lopes da Silva, 2006; Pfurtscheller & Neuper, 1997). In our study, responses were bimanual in the Implementation task and lateralized in the Memorization task. However, we reasoned that our test is independent of such difference, as before target onset responses from both hands had to be prepared in the two tasks.

## 3. Results

### 3.1 Behavioral results

Concerning behavioral performance, we expected valid retro-cues to have a beneficial effect on reaction times and error rates in both tasks (Souza & Oberauer, 2016). Repeated measures ANOVA on RTs confirmed a significant main effect of Load^3^ (*F*_1.44, 49.06_ = 104.65, *p* < 0.001, η^2^_p_ = 0.76): RTs increased with the number of mappings selected by the retro-cue (Load 1: *M* = 978ms, *SD* = 196, Load 2: *M* = 1168ms, *SD* = 150, Load 4: *M* = 1266ms, *SD* = 142). The main effect of Task was also significant (*F*_1, 34_ = 237.20, *p* < 0.001, η^2^_p_ = 0.87), with slower RTs in Memorization (*M* = 1355ms, *SD* = 185) than Implementation (*M* = 907ms, *SD* = 160). The interaction between the two factors also resulted to be significant (*F*_1.32, 44.88_ = 4.42, *p* = 0.031, η^2^_p_ = 0.11). More specifically, the effect of Load was larger in the Implementation task (*F*_1.44, 49.06_ = 93.20, *p* < 0.001) compared to the Memorization task (*F*_1.44, 49.06_ = 45.50, *p* < 0.001) (Figure 2a).

**Figure 2:**
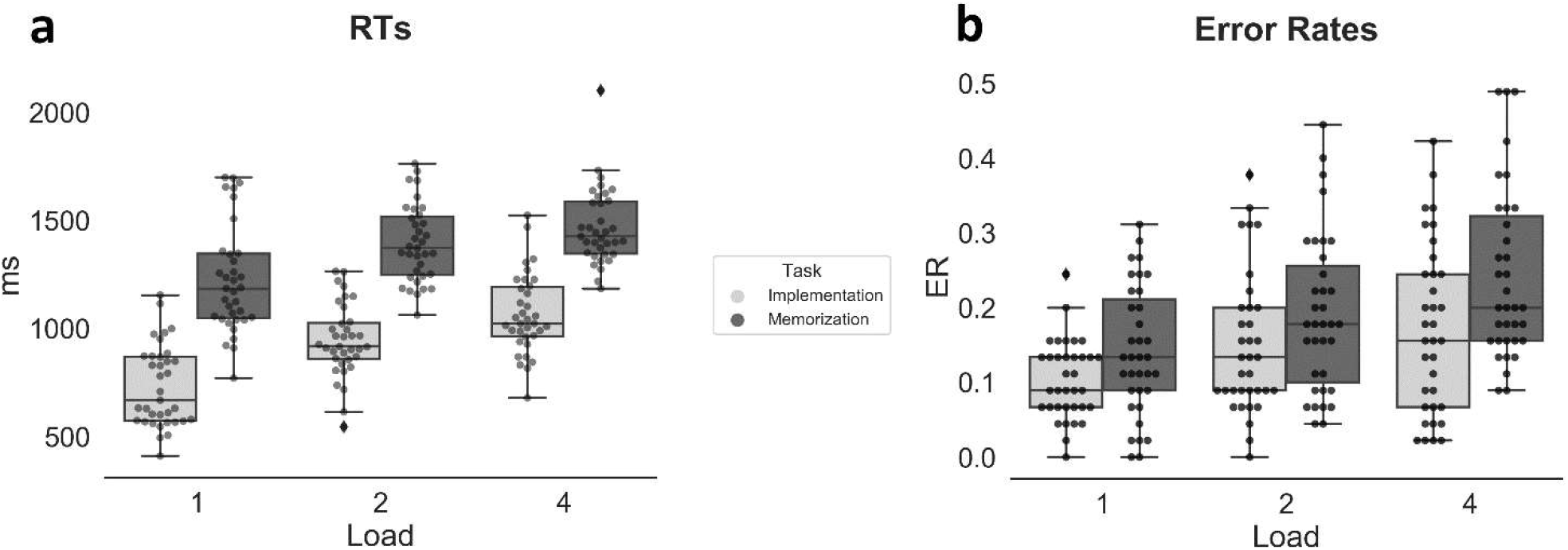
Behavioral results. **a**) Reaction times (ms). **b**) Error rates. Both measures confirmed a benefit in performance (i.e., faster RTs and less errors) when the Retro-Cue selected less mappings. In each boxplot, the thick line inside box plots depicts the second quartile (median) of the distribution (n = 35). The bounds of the boxes depict the first and third quartiles of the distribution. Whiskers denote the 1.5 interquartile range of the lower and upper quartile. Dots represent individual subjects’ scores.

Error rates showed a significant main effect of Task (*F*_1, 34_ = 11.08, *p* = 0.002, η^2^_p_ = 0.25) and of Load (*F*_1.67, 57.91_ = 25.65, *p* < 0.001, η^2^_p_ = 0.43). Participants were significantly more accurate in the Implementation task (*M* = 0.14, *SD* = 0.09) compared to the Memorization task (*M* = 0.19, *SD* = 0.10), and were more accurate when less items were selected by the retro-cue (Load 1: *M* = 0.12, *SD* = 0.07, Load 2: *M* = 0.17, *SD* = 0.10, Load 4: *M* = 0.20, *SD* = 0.12). The interaction between the two factors was not significant (*p* = 0.266) (Figure 2b).

Regarding catch trials, participants could successfully detect a new image in both the Memorization task (error rate in catch trials: *M* = 0.13, *SD* = 0.08) and the Implementation task (*M* = 0.09, *SD* = 0.07). Nevertheless, they were significantly less accurate (*t*_*34*_ = 3.16, *p* = 0.003, *d* = 0.53) in the Memorization task.

Such differences in performance between Implementation and Memorization are not surprising and can be at least partially attributed to the task-specific processes taking place after the occurrence of the target (Formica et al., 2020; Muhle-Karbe et al., 2017). Specifically, the Memorization task requires the comparison of the presented mapping to the encoded ones, the identification of the response side, and the execution of the intended response. On the contrary, the Implementation task only involves the identification of the target stimulus and the execution of the associated motor response.

## 3.2 EEG Results

### 3.2.1 Contralateral alpha power decrease

As hypothesized, we observed a stronger reduction in alpha power in contralateral compared to ipsilateral electrodes across loads 1 and 2 both in Implementation (*P* = 0.013, cluster-corrected) and Memorization (*P* = 0.003, cluster-corrected) tasks (see Supplementary material for the same analysis performed separately for the two load conditions). The significant clusters spanned the time window around 600 − 800ms. Notably, we did not find any difference between the two tasks (no cluster survived multiple comparison correction, even at a cluster threshold of *P* = 0.1), supporting our hypothesis that the mechanisms allowing for attentional orienting are engaged to a similar extent in the two tasks (Figure 3a). To qualitatively confirm our electrodes selection and the location of the significant effect, contralateral minus ipsilateral half-topographies are reported separately for each task (Figure 3b).

**Figure 3:**
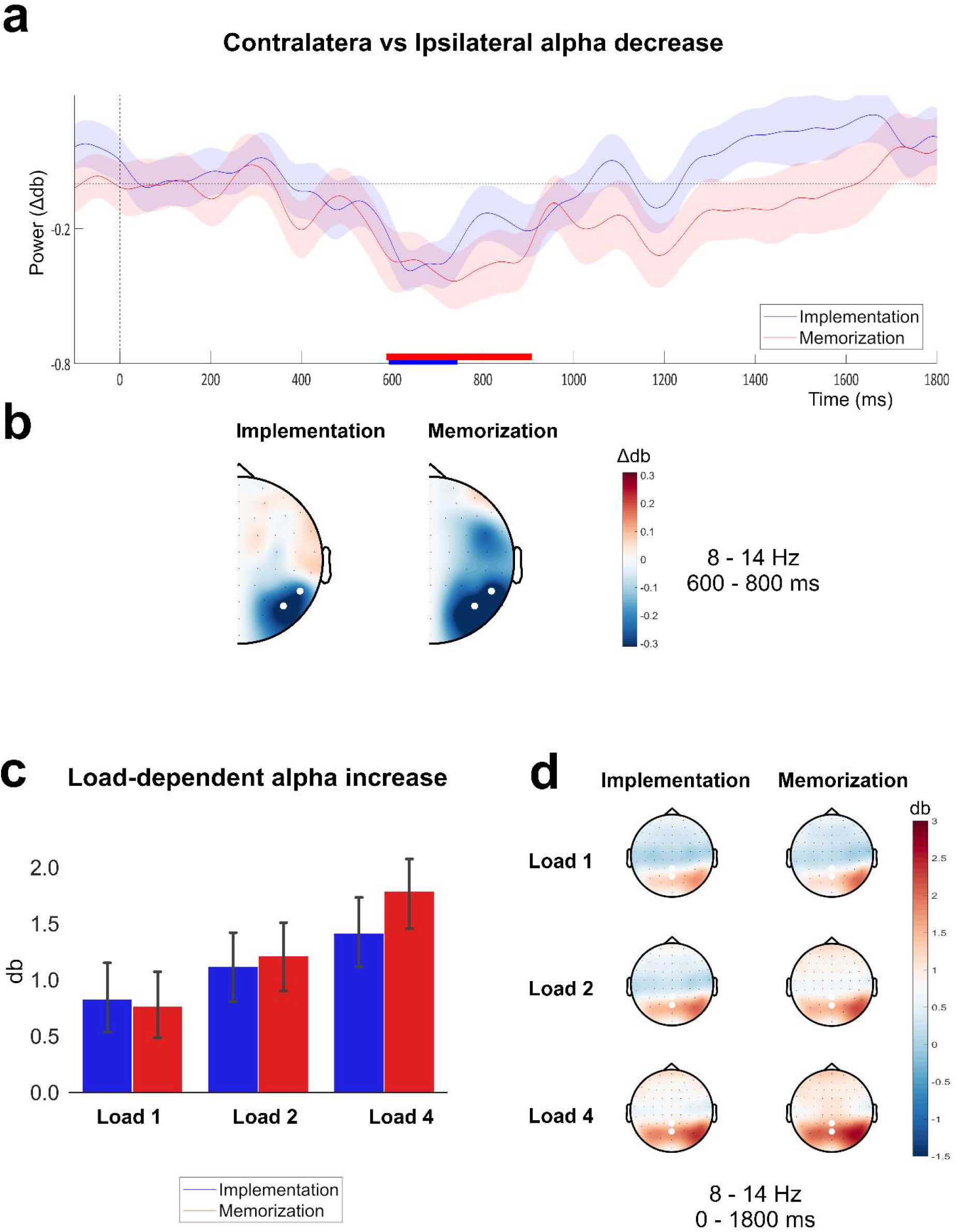
Alpha power dynamics. **a**) Time courses of the difference waves (contralateral minus ipsilateral time course) of alpha power activity between contralateral and ipsilateral electrode clusters (PO8 - P8 and PO7 - P7). Both difference waves show a significant deflection from 600 to 800ms after the onset of the retro-cue. Horizontal lines above the x-axis indicate significant temporal clusters (blue: Implementation, red: Memorization), obtained with cluster-based permutation testing). Shading represents the s.e.m., calculated across participants (n = 35). **b**) Half topographies showing differences in alpha power for contralateral minus ipsilateral electrodes collapsed across hemispheres. White dots indicate the a-priori selected electrodes used in the cluster-based permutation analysis. **c**) Alpha power values averaged in the time window 0 −1800 ms after Retro-Cue onset from electrodes Pz and POz. Error bars represents the s.e.m. calculated across participants (n = 35). **d**) Topographies of each Task x Load condition. White dots indicate the a-priori selected electrodes (Pz and POz) from which activity was extracted for the ANOVA.

### 3.2.2 Load-dependent alpha increase

We predicted alpha power to also track the number of items selected by the retro-cue to be relevant for the ongoing task. The repeated measures ANOVA on the averaged alpha power over Pz − POz in the whole CTI (0 − 1800ms) revealed a significant main effect of Load (*F*_2, 68_ = 24.81, *p* < 0.001, η^2^_p_ = 0.42) (Figure 3c). Specifically, alpha power values increased with the number of retained items (Load 1 vs Load 2: *t* = −3.24, *p* = 0.006 Bonferroni-corrected; Load 2 vs Load 4: *t* = −3.38, *p* < 0.001 Bonferroni-corrected), both in Implementation (Load 1: *M* = 0.83, *SD* = 1.83, Load 2: *M* = 1.12, *SD* = 1.83, Load 4: *M* = 1.42, *SD* = 1.92) and Memorization (Load 1: *M* = 0.77, *SD* = 1.78, Load 2: *M* = 1.22, *SD* = 1.87, Load 4: *M* = 1.79, *SD* = 1.90). Crucially, neither the effects of Task nor the interaction Task x Load were significant (*p*s > 0.1). Scalp topographies of alpha power across the entire CTI are reported separately for each Task and Load condition (Figure 3d).

### 3.2.3 Mid-frontal theta in Implementation

We expected to find stronger theta power over mid-frontal electrodes in the Implementation task compared to the Memorization task. Supporting this prediction, we found a significant cluster (P = 0.04, corrected for multiple comparisons) when comparing the power estimates of the two tasks in the frequency range 3 − 7 Hz from electrodes Fz and AFz (Figure 4). On the contrary, the test aimed at detecting a main effect of Load approached but did not reach significance (P = 0.09).

**Figure 4:**
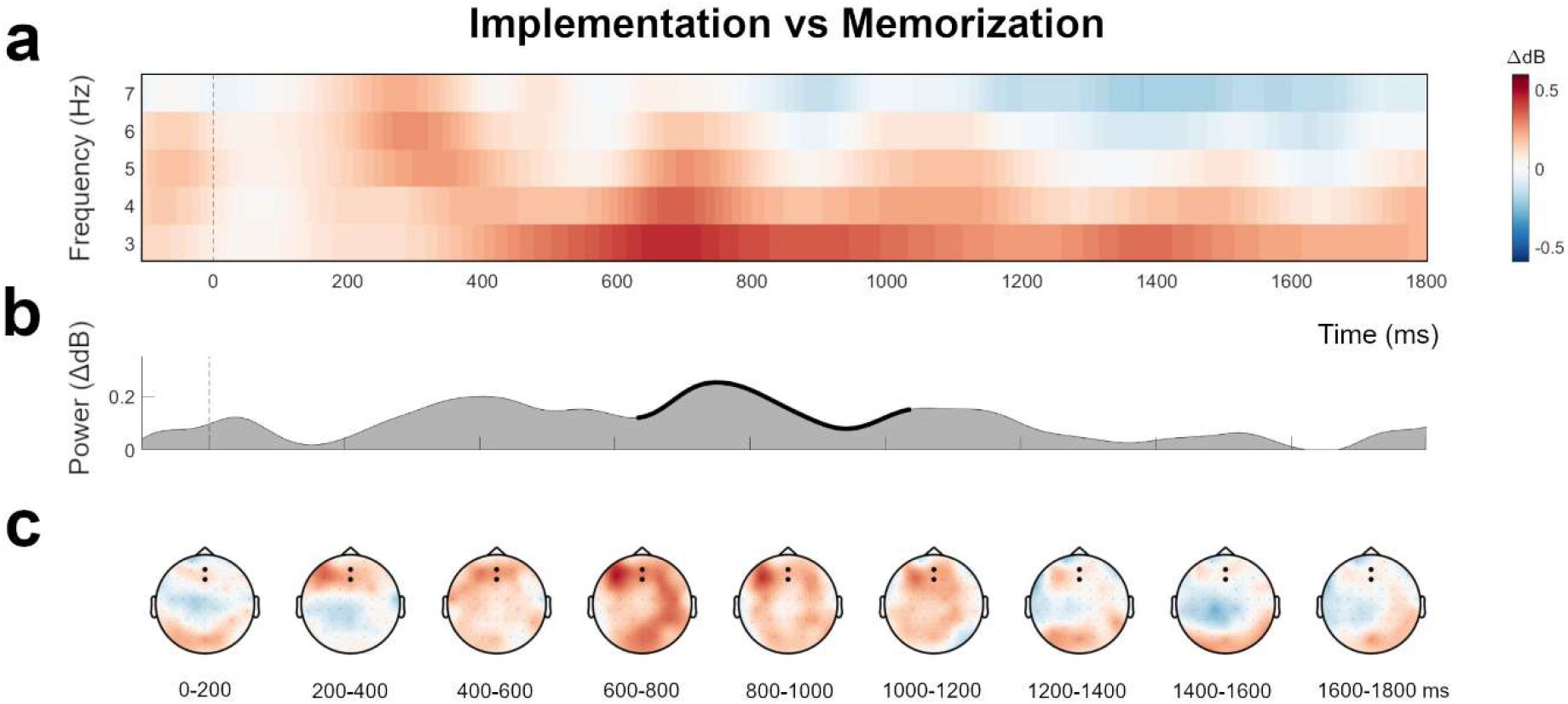
Theta dynamics in Implementation compared to Memorization. **a)** Time-frequency power spectrum extracted from the electrodes Fz and AFz. **b**) Averaged power time course (3 − 7 Hz) from the electrode Fz and AFz. Thicker outline indicates timepoints belonging to the significant cluster. **c)** Each topography represents the averaged activity in the frequency range 3 − 7 Hz, in 200 ms time windows within the CTI (0 − 1800 ms). Black dots indicate the electrodes used for the statistical analysis.

Moreover, our corollary hypothesis concerning theta dynamics predicted such effect to be modulated by the number of mappings to be implemented or memorized. Specifically, we expected that in Load 4 Implementation and Memorization would not differ, in line with behavioral findings showing that 4 novel instructions cannot be proceduralized at once and would therefore be maintained in a declarative format in both conditions (Liefooghe et al., 2012). However, the effect of the interaction resulted to be not significant (P = 0.53). Despite the lack of a significant interaction, and thus the impossibility to draw any strong conclusion from post-hoc tests, we explored the effect of Task across different Load levels by means of pairwise comparisons. We observed a significant cluster only in the contrast Implementation 2 vs Memorization 2 (P = 0.02), whereas no cluster was observed for Load 4, and no cluster survived correction in Load 1 (P = 0.23).

One potential confound when investigating activity in low frequency bands is that this might be reflecting sustained ERPs. To rule out this possibility we compared across tasks the Contingent Negative Variation (CNV), a slow negative potential over centro-frontal electrodes during preparation for an upcoming target (Walter, Cooper, Aldridge, McCallum, & Winter, 1964). The CNV did not show a differential pattern between Implementation and Memorization (see Supplementary material), suggesting that our frontal activity is better described by low frequency oscillations, rather than a sustained ERP. Furthermore, at the lowest estimated frequency of 2 Hz, the wavelet was two-cycles long, i.e. spanning a 1s interval, which is considered to be selective to true oscillations and able to disentangle them from non-oscillatory signal (Cohen, 2014a). Finally, interpreting this frontal activity as the correlate of target expectancy would not account for the effect specific to Load 2 observed in time-frequency analyses.

### 3.2.4 Motor-related mu and beta suppression in Implementation

We hypothesized that proceduralization would induce increased recruitment of motor and premotor regions, in preparation for the instructed upcoming movement, compared to Memorization. Specifically, we predicted a suppression of oscillations in the mu (8 − 12 Hz) and beta (15 − 30 Hz) frequency ranges, which have been shown to reflect motor cortex activity (Cheyne, 2013; Pineda, 2005; Rhodes et al., 2018; Schneider et al., 2017; Tzagarakis et al., 2015). To test for this hypothesis, we again used cluster-based permutations and we tested the effects of Task, Load, and their interaction, separately for each frequency range of interest (i.e. mu and beta). In line with our prediction, we found significant clusters when contrasting the two Tasks both in the mu (P = 0.01) and beta (P = 0.03) frequency ranges. This finding supports the idea that proceduralization involves a recruitment of motor areas for the activation of the instructed motor plan (Figure 5).

**Figure 5:**
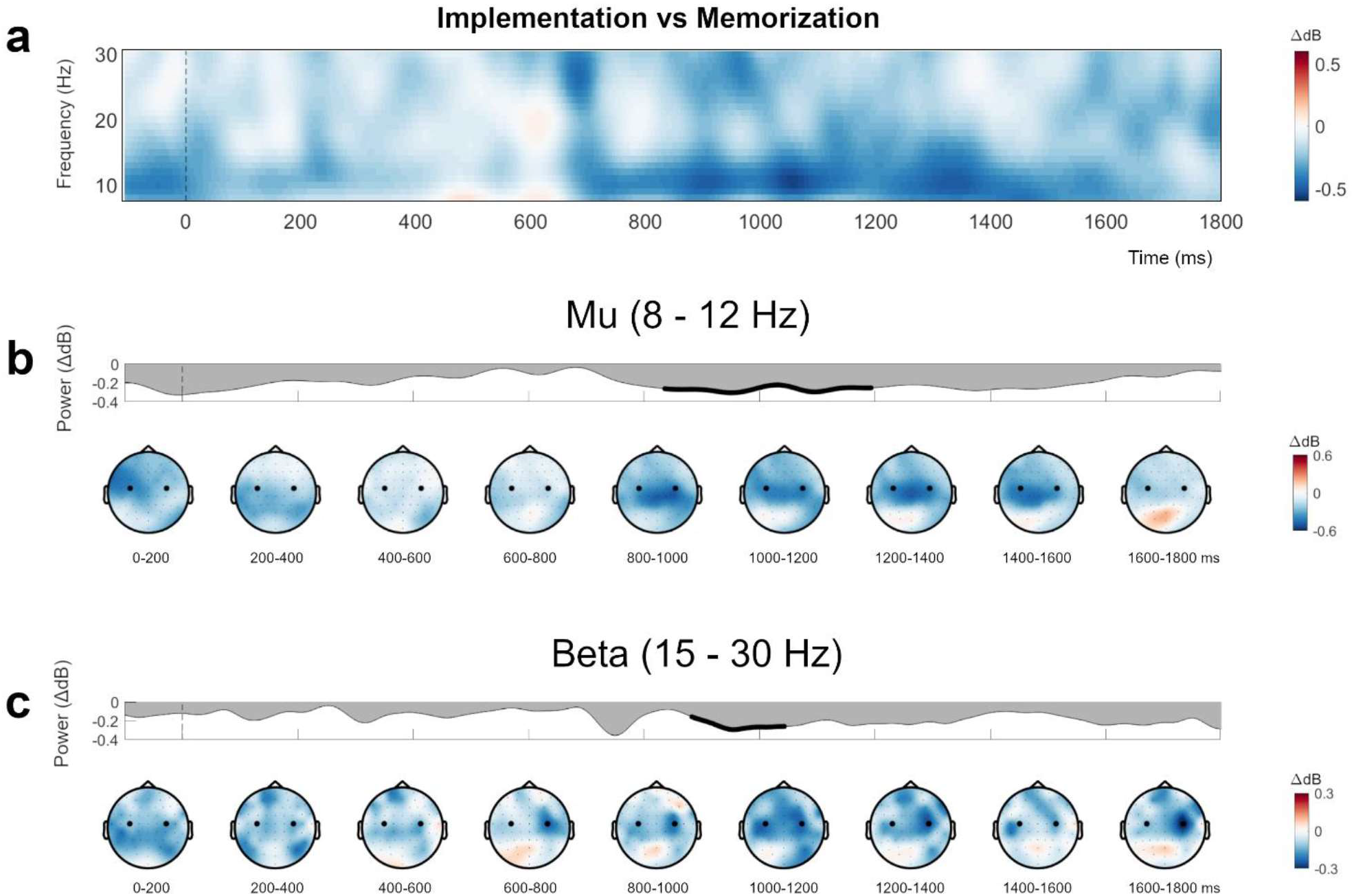
Mu and Beta dynamics in Implementation compared to Memorization. **a)** Time-frequency power spectrum from electrodes C3 and C4. **b)** Averaged power time course in the Mu (8 − 12 Hz) frequency range from electrodes C3 and C4: thicker outline indicates timepoints belonging to the significant cluster. Topographies in 200 ms time windows: black dots indicate the electrodes used for statistical analyses. **c)** Averaged power time course in the Beta (15 − 30 Hz) frequency range from electrodes C3 and C4: thicker outline indicates timepoints belonging to the significant cluster. Topographies in 200 ms time windows: black dots indicate the electrodes used for statistical analyses.

Similarly, the effect of Load resulted to be significant in both mu (P < 0.001) and beta (P < 0.001) range, suggesting that these motor-related oscillatory features are also sensitive to the number of items being handled.

As for theta, our secondary hypothesis was that the differences in mu and beta dynamics between tasks would be modulated by the number of mappings. This hypothesis would be in line with a capacity limit in the amount of instructions that can be proceduralized at once, and thus converted in an action plan. However, in neither of the two frequency bands the cluster-based permutations tests revealed significant clusters for an effect of such interaction (mu: P = 0.14, beta: P = 0.57). As exploratory follow-up analyses we performed pairwise comparisons separately for each Load level (i.e., Implementation 1 vs Memorization 1, Implementation 2 vs Memorization 2, Implementation 4 vs Memorization 4). Although interpretations of these results have to be cautious and do not allow to draw any strong statistical conclusion, it is interesting to observe that again only in Load 2 the two tasks showed clusters of different activity (beta: P1 < 0.001, P2 = 0.02, P3 = 0. 21, mu: P1 = 0.003, P2 = 0.006, P3 = 0.042). On the contrary, comparisons in Load 1 (beta: P = 0.19, mu: no cluster) and Load 4 (beta: P = 0.35, mu: P = 0.13) yielded no significant clusters.

## 4. Discussion

In the present study, we investigated the spectral and temporal dynamics underlying novel instruction implementation. In line with our predictions and the framework proposed by Myers and colleagues (2017), we observed an analogous unfolding of early attentional mechanisms across tasks, reflected in similar patterns of alpha power dynamics. Crucially, Implementation and Memorization also showed differences in frontal theta, and central mu and beta oscillations, suggesting that Implementation-specific processes go beyond the attentional prioritization of the relevant items. These mechanisms are likely involved in the reformatting of the selected S-R mappings into a behavior-optimized, action-bound process requiring the exertion of cognitive control and resulting in the preparation of the instructed motor plan (van Ede, 2020).

An underlying assumption of our study is that the intention to implement the instructed mappings is a crucial necessary factor to trigger the reformatting from declarative to procedural state (Liefooghe et al., 2012). This exclusive role of the intention to implement has been questioned by some recent behavioral evidence, suggesting an automatic reformatting of the instruction independent of task demands (Liefooghe & De Houwer, 2017). This might be due to the response set used in the specific task, or to other idiosyncratic factors. Despite these inconsistencies in the behavioral measures, the dissociation between Implementation and Memorization is well established at the neural level (Demanet et al., 2016; González-García et al., 2019; Muhle-Karbe et al., 2017). Here, we show such differences are reflected in oscillatory mechanisms.

### 4.1 Alpha-mediated attentional orienting is analogous in Implementation and Memorization

Activity in the alpha frequency band has been consistently associated with attentional processing (van Ede, 2017). In particular, posterior alpha power decreases over posterior electrodes contralateral to the attended hemifield, both in perceptual and in WM tasks (Jensen & Mazaheri, 2010; Poch et al., 2017; Sauseng et al., 2005). In our experiment, participants were encouraged to focus on the selected items and discard the unselected ones. Coherently, in both tasks we found after the retro-cue a significant suppression of alpha oscillations contralateral to the attended hemifield. The most influential model proposed to account for this phenomenon is referred to as *Gating by Inhibition* (Jensen & Mazaheri, 2010). According to this framework, top-down alpha modulation of sensory cortices allows for the inhibition of irrelevant inputs, contributing to the creation of a functional network optimized to perform the task (Mazaheri et al., 2014; Van Diepen, Foxe, & Mazaheri, 2019). Our clusters of alpha lateralization extended around 600 − 800ms after retro-cue. This dynamic is consistent with previous findings on lateralized retrospective orienting of internal attention, showing modulations within 500 and 1000ms (Mok et al., 2016; Poch et al., 2017, 2018; Wallis et al., 2015; Wolff, Ding, Myers, & Stokes, 2015; Wolff, Jochim, Akyürek, & Stokes, 2017). The occurrence of the selection process is further supported by load-dependent centro-posterior alpha power modulation. The amplitude of alpha power has been observed to be sensitive to the number of items retained during a WM task (Fukuda et al., 2015; Jensen, 2002). In both our tasks, alpha increased with load. Crucially, we observed no task-dependent differences in the dynamics of alpha oscillations, suggesting that the information provided by the retro-cue is used to analogously orient attention and select the items that are likely to be probed, independently of task demands.

### 4.2 Low frequencies oscillations for S-R binding in Implementation

We predicted low frequency oscillations over frontal sensors to be associated with the proactive reformatting of the S-R mapping(s) into an action-oriented code. The crucial contribution of the PFC in quickly converting symbolic instructions into task-sets for prospective action has been widely established in several fMRI studies (Brass, Wenke, Spengler, & Waszak, 2009; Cole, Bagic, Kass, & Schneider, 2010; Dumontheil, Thompson, & Duncan, 2011; González-García et al., 2017; Hartstra et al., 2011; Palenciano, González-García, Arco, & Ruz, 2019; Ruge & Wolfensteller, 2010). In particular, declarative maintenance and implementation of novel mappings can be dissociated using univariate and multivariate analyses in prefrontal regions, such as the inferior frontal junction (IFJ) and the inferior frontal sulcus (IFS) (Bourguignon et al., 2018; Demanet et al., 2016; Hartstra et al., 2011; Muhle-Karbe et al., 2017). Activation of these areas has been interpreted as reflecting the creation and the activation of procedural condition-action rules from the instructed mappings, a process that is thought to involve the manipulation of the available information and the exertion of cognitive control (Brass et al., 2017). Analogously, the need for top-down control over complex goal-directed cognitive operations is traditionally linked with oscillations in the theta frequency range over prefrontal cortices (Cavanagh & Frank, 2014; Cooper et al., 2019). Moreover, recent influential models of large-scale brain interactions attribute to theta the crucial role of orchestrating information exchange between distant areas by synchronizing their firing pattern (Fries, 2005, 2015; Lisman & Jensen, 2013; McLelland & VanRullen, 2016; Verguts, 2017). We hypothesized the Implementation task in our experiment to require a more extensive allocation of cognitive control, given the need to manipulate the declarative representations of the selected S-R mappings into their procedural counterparts, leading to increased power in low frequencies over frontal sensors. Supporting this prediction, our Implementation and Memorization tasks differed in the extent to which they engaged oscillations in the theta frequency range over prefrontal regions.

Previous studies show mixed findings with respect to the role of midfrontal theta power in dissociating maintenance and manipulation of WM items. Berger and colleagues (2019) reported no differences in theta power between retention and mental spatial mirroring (i.e., manipulation) of a grid of colored dots. On the contrary, the study by Itthipuripat and colleagues (2013) used a sequence updating paradigm and revealed increased theta for both contrasts Manipulation vs Maintenance and Manipulation (correct trials) vs Manipulation (incorrect trials). Interestingly, they found low theta (2 − 4 Hz) to be sensitive to manipulation demands, whereas high theta (5 − 7 Hz) increased in both contrasts (Itthipuripat et al., 2013). Although the exact boundaries of our significant cluster cannot be strongly interpreted due to the nature of the statistical approach we adopted (see statistical Methods section), the effect of Task we find in our study appears to be predominant in the lower frequencies. One possibility is that different frequencies within the theta band serve different cognitive functions. This view has recently been gaining support. Evidence in favor of a more fine-grained characterization of theta oscillations include variations in theta peak frequency depending on task demands and difficulty (Senoussi et al., 2020), and differences in the functional significance of high (posterior) and low (anterior) hippocampal theta (Goyal et al., 2020).

A functional role has been attributed to theta oscillations in the prioritization of relevant information in visual WM. Oscillations in the delta/low theta (2 − 6 Hz) range have been observed during switches of internal attention from one search template to another for prospective use (de Vries et al., 2018; de Vries, van Driel, & Olivers, 2019b; Senoussi et al., 2019). Such involvement has also been causally tested by applying rhythmic (5 Hz) and arrhythmic transcranial magnetic stimulation to the lateral PFC during a change detection task with retro-cues (Riddle et al., 2020). Rhythmic stimulation caused an entrainment of the ongoing theta oscillations and benefitted performance, compared with the arrhythmic train of pulses (Lakatos, Gross, & Thut, 2019; Riddle et al., 2020; Thut, Schyns, & Gross, 2011; Thut, Veniero, et al., 2011).

In our study, the difference in theta activity observed between tasks suggests that its function might go beyond the exertion of short-lasting top-down control over attentional prioritization, but also encompasses the reformatting of the prioritized memoranda into behavior-guiding representations. This is possible for the Implementation task, whereas in the Memorization task the prioritized representations can only be used as a template to guide the upcoming task (de Vries, Slagter, & Olivers, 2019; Olivers & Eimer, 2011; Olivers, Peters, Houtkamp, & Roelfsema, 2011).

Therefore, the stronger theta activity in Implementation likely reflects the inherently more extensive reformatting process of proceduralization, which we assume involves a theta-driven binding between the stimulus and its corresponding motor plan (Combrisson et al., 2017; Engel, Roelfsema, Fries, Brecht, & Singer, 1997; Verguts, 2017). The Memorization task also requires the prioritization and the binding of stimulus and response, but this would engage theta dynamics to a lesser extent, given the impossibility to activate a motor plan.

Although strong conclusions cannot be made with respect to timing due to the temporal smearing inherent to time-frequency analyses, attentional orienting and theta-mediated binding seem to be occurring simultaneously. The temporal extents of the two clusters are, in fact, largely overlapping. Therefore, our results hint at a parallel engagement of the processes that underlie these frequencies.

### 4.3 Motor preparation in Implementation is reflected in mu and beta suppression

Previous fMRI studies showed that the proactive proceduralization of novel instructions is accompanied by increased neural activity in pre-motor areas, suggesting motor preparation (Bourguignon et al., 2018; Hartstra et al., 2011, 2012; Muhle-Karbe et al., 2017; Ruge & Wolfensteller, 2010). In current views, the recruitment of motor areas is thought to boost instruction implementation by enabling mental simulation of the overt application of the instructed mappings (Brass et al., 2017). Motor imagery has been associated with suppression in mu and beta bands over motor cortices (Cheyne, 2013; McFarland et al., 2000; Pfurtscheller et al., 2006; Pfurtscheller & Neuper, 1997; Pineda, 2005), which we therefore predicted to occur while the participant is preparing to implement a specific motor plan, as opposed to the declarative maintenance of the mapping. Moreover, the engagement of motor areas during implementation is consistent with the creation of a functional network optimized for the rapid and efficient execution of the mappings.

In line with our hypothesis, we found these markers of motor preparation to be higher in Implementation compared to Memorization, already during the CTI. While in both tasks participants had to perform an overt motor response at the end of the trial, only during Implementation they could proactively start preparing their response to the upcoming target. This rules out the possibility that the observed differences reflect more general mechanisms of unspecific motor energization, temporal estimation or target expectancy (Nobre & Van Ede, 2018; Van Elswijk, Kleine, Overeem, & Stegeman, 2007; Wiener, Parikh, Krakow, & Coslett, 2018). Overall, our results suggest that preparing to execute novel mappings involves the activation of a specific motor plan corresponding to the instructed response, to a much larger extent then the motor engagement observed in Memorization.

Concerning the timing of the implementation-induced motor engagement, the temporal extent of the significant cluster suggests a later onset with respect to alpha and theta dynamics. Although strong conclusions cannot be drawn, based on this interpretation of our permutation results and theoretical models such as Myers’ (2017), we are inclined to speculate that motor preparation is initiated following attentional selection of the relevant mappings. From this perspective, theta would act as an overarching mechanism to bind and control the task-optimized representations. Future studies should investigate the exact timing properties and the interplay between these processes.

### 4.4 Capacity limits of proceduralization

As a corollary hypothesis, we predicted that the capacity limits of proceduralization observed in behavioral studies (Liefooghe et al., 2012) would be reflected in stronger proceduralization-related oscillatory features with fewer mappings. However, we did not observe a significant interaction of Task and Load in none of our frequency bands of interest. It is possible that our design was not powered enough to detect such effects. Despite the not significant interaction, for exploratory purposes we looked at the effect of Task separately for each Load. Interestingly, we observed task-related differences only in Load 2. The interpretation of these results requires caution and can only be speculative given the exploratory nature of these analyses. Yet, they could reflect that relevant oscillations become crucial when in need to coordinate multiple bindings, but still within capacity limitations. From this perspective, Load 4 would show no differences because too many S-R mappings cannot be simultaneously recoded into action-oriented representations (Liefooghe et al., 2012), and Load 1 would instead allow a more ballistic implementation of the prepared motor plan, without the need to carefully coordinate different competing procedural representations, as is the case in Load 2. These exploratory results pave the way for future research investigating more accurately the capacity limits of proceduralization: how they are overcome with extensive practice and how they reflected in power modulations.

## 5. Conclusions

Our study shed light for the first time on the oscillatory dynamics associated with the retrospective prioritization and proceduralization of novel instructions. We showed that alpha-mediated mechanisms of attentional orienting are in place to prioritize the relevant items, independently from the upcoming task demands. Conversely, other neural features were sensitive to the task. The implementation of novel mappings, as opposed to their declarative maintenance, is characterized by increased power in low frequencies and by stronger suppression of mu and beta activity. The former is consistent with the purported role of theta oscillations in cognitive control for the successful binding of stimulus and response into a behavior-optimized procedure, and the latter to reflect motor preparation of the instructed motor responses. Overall, our results support the idea that under optimal task conditions, proceduralization can occur proactively in preparation for the upcoming response execution demands.

Future research should investigate how proceduralization affects the underlying neural representation of the instructed mapping, under the assumption that it is recoded from a declarative to a procedural format, and how prefrontal regions mediate this reformatting process. Moreover, future studies should explore whether capacity limits of proceduralization (Liefooghe et al., 2012) are caused by concurrent activation of interfering motor plans or intrinsic to the neural mechanisms supporting the recoding of instructions into procedures, such as the theta-mediated binding.

## Acknowledgements

S.F. and C.G.G. were supported by Special Research Fund of Ghent University BOF.GOA.2017.0002.03. C.G.G. was additionally supported by the European Union’s Horizon 2020 research and innovation programme under the Marie Sklodowska-Curie grant agreement no. 835767. M.S. was supported by Research Foundation Flanders (FWO) grant number G012816N. MB is funded by an Einstein Strategic Professorship of the Einstein Foundation Berlin.

## Data and code availability

Data and code will be made available upon publication.

## Footnotes

1 The physical properties of the retro-cue could potentially produce a combination of exogenous and endogenous orienting of attention in Load 1 and 2 (i.e., one side of it is colored and thus more salient). However, we are not strongly interested in this distinction and aim at using the retro-cue as the event to locate in time the onset of different demands between tasks.

2 Electrodes selection for analyses in the mu and beta frequency ranges was based on the literature and on the clearer assumptions on the brain areas causing the expected oscillations (i.e., pre-SMA and M1). Moreover, we chose to not rely on an independent electrode selection as we did for theta because the FOOOF toolbox results in higher frequencies are more susceptible to muscular artifacts and therefore less accurate.

3 Mauchly’s test revealed that the assumption of sphericity is violated (p<0.05).Greenhouse-Geisser correction is applied here and in all results where the sphericity assumption is violated.

