## Supplementary Material for "Neural oscillations track the maintenance and proceduralization of novel instructions"

1. *FOOOF electrode selection*

As a first step, we computed the power spectra from all trials, independently of Task and Load conditions, for each participants and averaged across them.

2. *Contingent Negative Variation (CNV)*

The CNV is a well-established ERP component occurring between the presentation of a cue and the target. It consists of sustained negativity over central scalp electrodes. We tested whether the difference we found in theta between Implementation and Memorization could be better explained by this ERP signature of temporal expectation. We extracted the averaged ERP form the channel Fz, separately for each tasks and Load condition, baseline-correct to the average of the time window from -200 to 0 ms before the retro-cue onset. We used the cluster-based permutation approach (N permutations = 10000) to compare the two time-courses separately for each Load condition. In Load 1 and 2 we found no cluster of significant differences (see Figure S1). In Load 4, we observed a small significant cluster, but this is reflecting an early difference between the two Tasks in the post Retro-cue processing, not compatible with the expected CNV. Therefore, we concluded that our effect is better captured by low frequency oscillations rather than the sustained ERP.

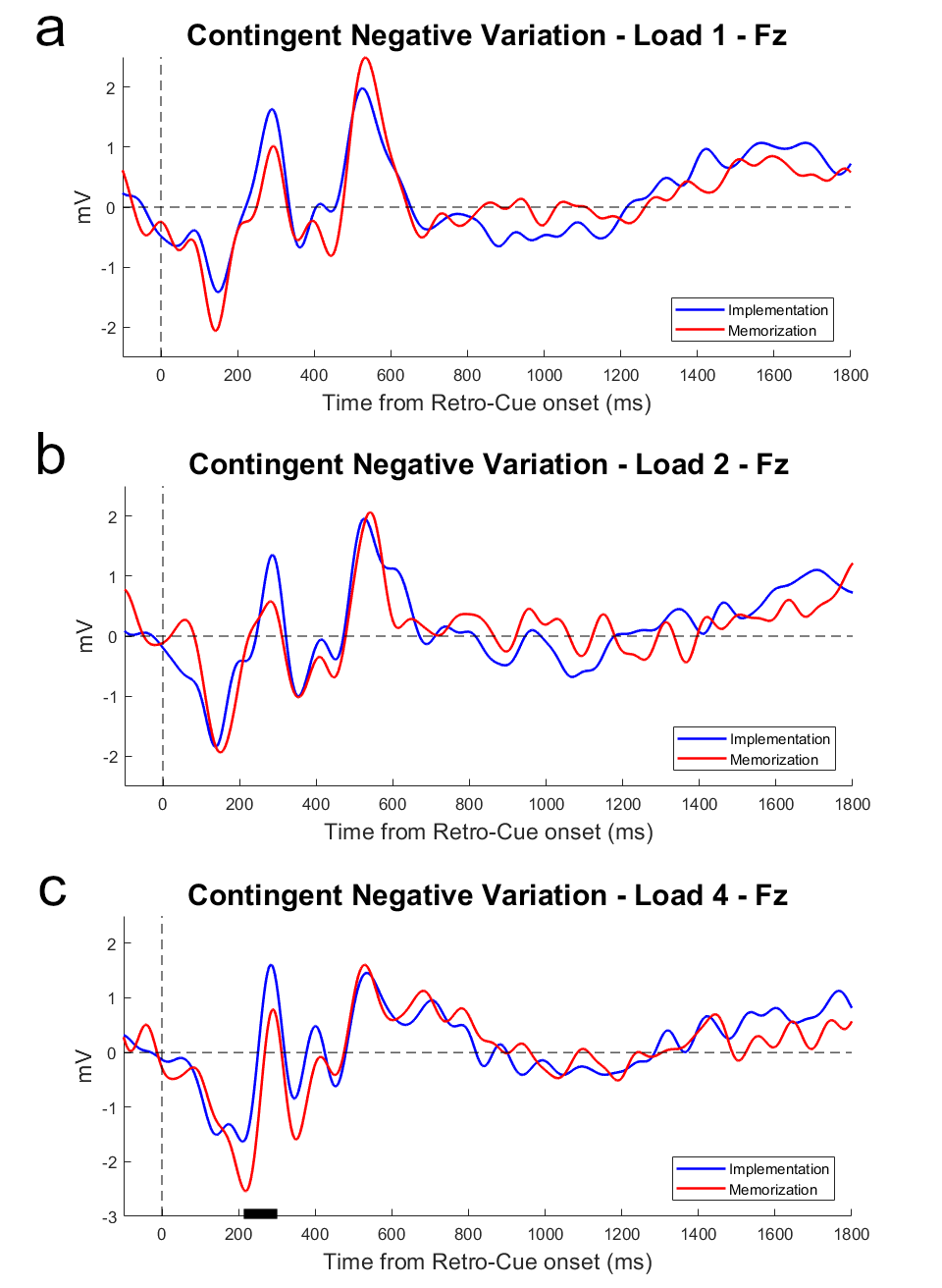

**Figure S2: Contingent Negative Variation**. ERPs averaged over the electrodes Fz, separately for Implementation and Memorization, for each Load condition (**a** – Load 1; **b** – Load 2; **c** – Load 4). For visualization purposes, the ERPs are filtered at 15 Hz. No differences were found between the two tasks.

3. *Contralateral alpha suppression – Load 1 and Load 2*

We performed a comparison between contralateral and ipsilateral alpha power time courses separately for Load 1 and Load 2. For Load 1, we didn’t find any significant cluster of activity, even though the scalp topographies in the time window of interest 600 – 800ms qualitatively show a slight suppression of alpha in the expected location. For Load 2, a significant cluster resisted correction for multiple comparisons in the Implementation task (*P* < 0.001, cluster-corrected), but no cluster resulted to be significant in the Memorization task (Figure S2).

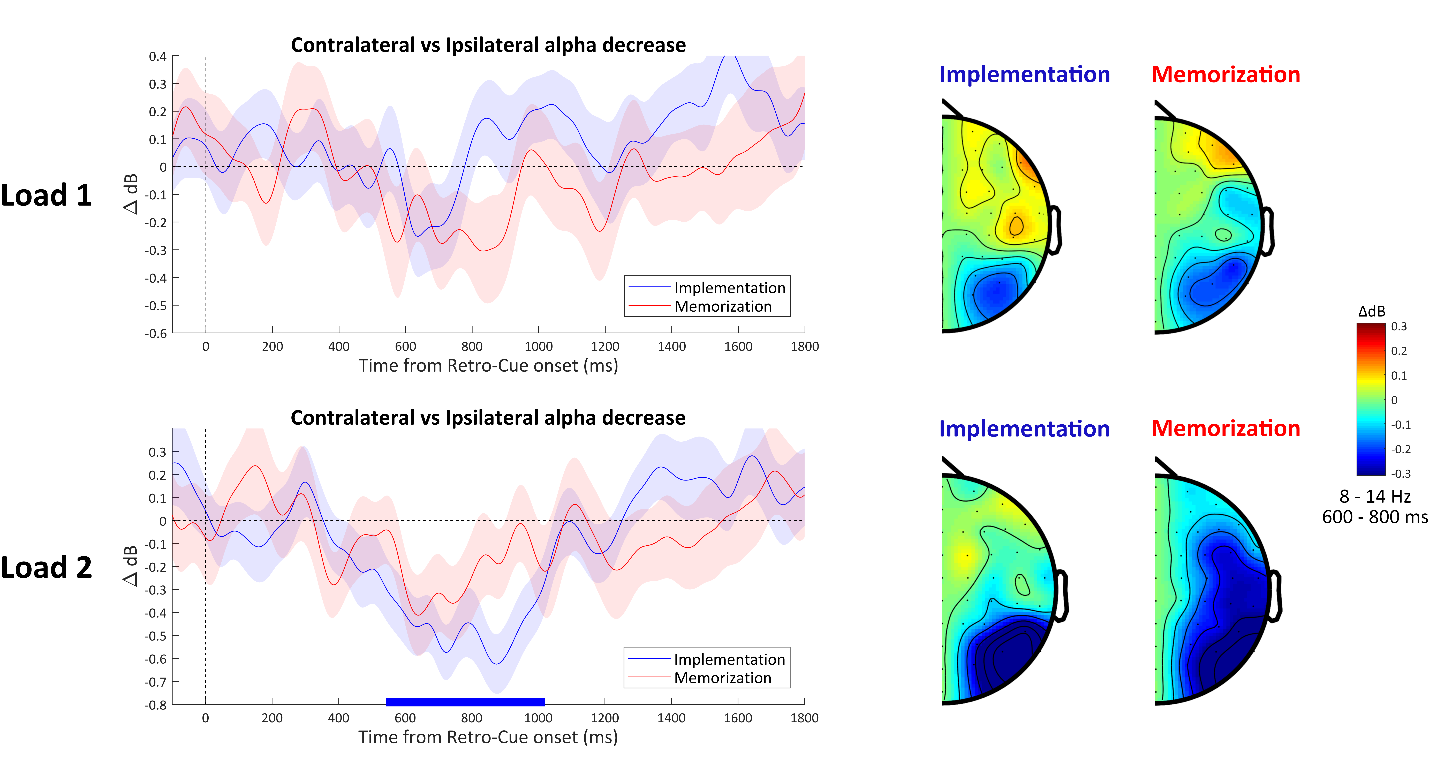

**Figure S2: Contralateral alpha decrease, separately by Load**. Time courses of the difference waves (contralateral minus ipsilateral time course) of alpha power activity between contralateral and ipsilateral electrode clusters (PO8 - P8 and PO7 - P7), separately for Load 1 and Load 2. Only Load 2 within the Implementation task shows a significant deflection from 600 to 1000ms after the onset of the retro-cue. Horizontal lines above the x-axis indicate significant temporal clusters (blue: Implementation, red: Memorization), obtained with cluster-based permutation testing). Shading represents ± 1 s.e.m., calculated across participants (n = 35). Half topographies showing differences in alpha power for contralateral minus ipsilateral electrodes collapsed across hemispheres.

To further test for differences between the two load conditions, we compared the difference waves for Load 1 and for Load 2 (averaged across tasks). The goal of this analysis was to capture differences in the time course of lateralized alpha power between loads. No cluster survived the correction for multiple comparisons. At the uncorrected significance level, the analysis showed differences between the two load conditions around the time window 700 – 1000ms, suggesting stronger suppression for Load 2 (Figure S3).

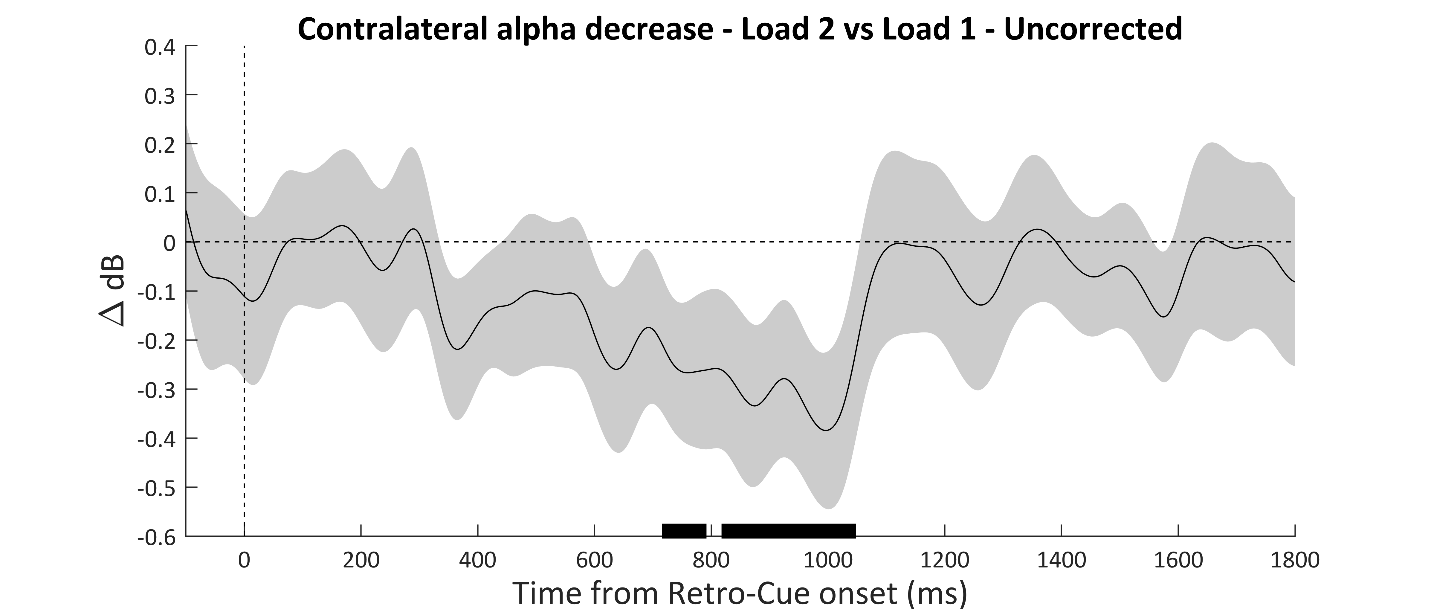

**Figure S3: Comparison of contralateral alpha decrease between Load 2 and Load 1, averaged across Tasks**. Time course of the difference waves (contralateral minus ipsilateral time course) of alpha power activity between contralateral and ipsilateral electrode clusters (PO8 - P8 and PO7 - P7). Difference waves were averaged across Tasks separately for Loads, and then Load 1 was subtracted to Load 2. The difference (uncorrected) suggests stronger suppression for Load 2 compared to Load 1.
